# Propionate metabolism dysregulation promotes drug-tolerant persister cell survival in non-small cell lung cancer

**DOI:** 10.1101/2025.10.15.682658

**Authors:** Bobak Parang, Liron Yoffe, Rabia Khan, Zhongchi Li, Michal J. Nagiec, Eric E. Gardner, Yiwey Shieh, Rachel S. Heise, Ashish Saxena, Nasser Altorki, John Blenis

## Abstract

Recent studies show that genetic sequencing can not fully explain drug resistance in non-small cell lung cancer (NSCLC), suggesting undiscovered non-genetic mechanisms that can enable cancer cell survival. Propionate metabolism is the pathway by which odd-chain fatty acids, branched chain amino acids, and cholesterol are metabolized. We have previously shown that methylmalonic acid (MMA), a byproduct of propionate metabolism that accumulates when the pathway is disrupted, can activate epithelial-to-mesenchymal transition (EMT) in cell lines. But the clinical significance of propionate metabolism in cancer patients is not known. Here we show, for the first time, that propionate metabolism is dysregulated in patients with non-small cell lung cancer. MMA is elevated in lung tumors and in the serum of patients with metastatic NSCLC. Metabolism of cobalamin associated B (MMAB), a key regulatory gene of propionate metabolism, is downregulated in NSCLC and drug-tolerant persister cells, leading to MMA accumulation and EMT activation. We show that restoring expression of MMAB in NSCLC enhances targeted therapy and suppresses TGFB signaling. These findings reveal propionate metabolism dysregulation as a non-genetic mechanism of drug resistance and highlight propionate metabolism as a potential therapeutic target.

## Introduction

Non-small cell lung cancer is the leading cause of cancer-related deaths worldwide (1). Epidermal growth factor receptor (**EGFR**)-mutant NSCLC is a subset of lung cancer driven by oncogenic mutations that render EGFR constitutively active. EGFR-mutant NSCLC kills approximately 250,000 patients worldwide annually (2, 3). The development of small molecule inhibitors, like osimertinib, that inhibit the EGFR kinase has dramatically improved outcomes for patients with metastatic disease(4), but invariably, residual disease persists and eventually gives rise to proliferative, resistant disease. These residual cancer cells are known as “drug tolerant persister” cells (**DTPs**) and are characterized by slow-cycling and activation of epithelial-to-mesenchymal transition (**EMT**) and injury-response transcriptional signatures (5). Enrichment of TGFβ, YAP, and β-catenin signaling have been implicated in sustaining DTP survival (6–8). Although our understanding of DTP transcriptional states has advanced significantly, our understanding of DTPs metabolic regulation remains incomplete. Indeed, a more complete understanding of DTPs metabolic states may provide new insights into vulnerabilities that could be targeted.

Propionate metabolism is the metabolic pathway by which odd-chain fatty acids, branched chain amino acids, and cholesterol proceed through a series of enzymatic reactions to produce succinate (9). Succinate then enters the TCA cycle to produce ATP. Appropriate regulation of propionate metabolism involves a complex set of proteins including propionyl-coA carboxylase subunit alpha (**PCCA**), propionyl-coA carboxylase subunit beta (**PCCB**), methylmalonyl coA mutase (**MUT**), metabolism of cobalamin associated b (**MMAB**), metabolism of cobalamin associated a (**MMAA**), and methylmalonyl coA epimerase (**MCEE**). Inactivating mutations or loss of expression of *MUT, MMAB, MMAA*, or *MCEE* can disrupt the pathway and lead to accumulation of the metabolic by product methylmalonic acid (**MMA**)(10–12). It is known that pathologic germline mutations in these enzymes can obstruct propionate metabolism, and as a result, MMA levels accumulate. These inborn errors of metabolism are collectively referred to as “methylmalonic acidemias.” Unfortunately, these alterations lead to severe complications including death (13). A deficiency in adenosylcobalamin (vitamin B12), a co-factor for the enzyme MUT, can also slow the pathway and lead to a rise in serum MMA.

Work over the last decade has revealed that errors in propionate metabolism are relevant to more than just methylmalonic acidemias and vitamin B12 deficiency. Population level studies have shown that MMA levels increase in the serum with age (14–17) while controlling for kidney function and vitamin B12 levels. Furthermore, MMA levels correlate with mortality: higher MMA levels associate with greater overall mortality (16, 17). Recently, our group has shown that MMA can activate EMT in lung cancer cell lines and induce cancer-associated fibroblast (**CAF**) formation from fibroblasts derived from the lung (18, 19), promoting drug resistance.

Whether MMA levels are elevated in cancer patients is unknown. More broadly, propionate metabolism has not been well characterized across various diseases. We hypothesized that propionate metabolism is dysregulated in non-small cell lung cancer. In this study, we show for the first time that propionate metabolism dysregulation is a feature of NSCLC and can drive DTP survival. Collectively, this study provides another layer of complexity around non-genetic mechanisms of acquired drug resistance that limit the efficacy of targeted therapy in lung cancer.

## Results

### Methylmalonic acid (MMA) is elevated in lung cancers and patients

The hallmark of propionate metabolism dysregulation is accumulation of MMA. To test if MMA levels were elevated in NSCLC, we performed targeted polar metabolomics on resected tumors with paired adjacent non-malignant tissue from each patient. To rule out the confounding effects of kidney dysfunction and vitamin B12 deficiency, we performed metabolomics on specimens from patients with both normal vitamin B12 levels (defined as >300 pg/dl) and renal function (defined as eGFR >60). Our data showed that MMA was elevated approximately 2-fold in tumor tissue compared to paired adjacent non-malignant tissue within the lung (**Figure 1A**). MMA, succinate, and lactate were the most significantly enriched metabolites in tumor tissue compared to non-malignant lung tissue (**Supplemental Figure 1**).

**Figure 1.**
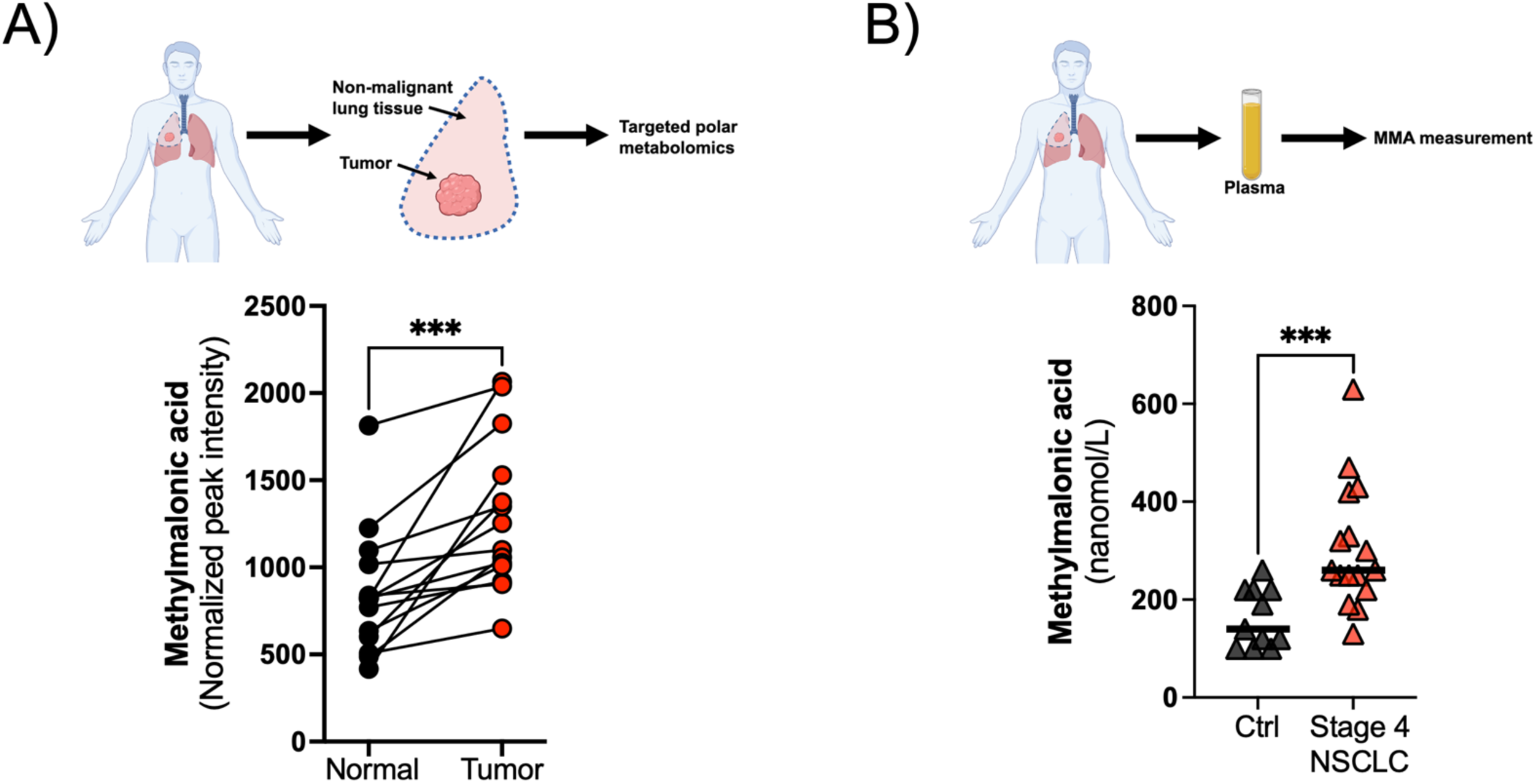
Methylmalonic acid is elevated in the tumors and serum of patents with non-small cell lung cancer. **(A)** Targeted metabolomics of resected non-small cell lung cancer and matched non-malignant tissue (n=12). Paired Student’s t-test. ***p<0.001 **(B)** Serum MMA levels from patients with treatment naïve, Stage 4 NSCLC compared to patients who had a history of stage 1 or 2 lung cancer and are cancer free. Control (n=11), Stage 4 (n=16). Unpaired Student’s t-test ***p<0.001.

To test if *serum* MMA is elevated in patients with metastatic NSCLC, we prospectively measured MMA levels in patients with treatment-naïve, stage 4 NSCLC. As a control, we measured serum MMA in patients who had a history of early-stage NSCLC and were disease free after surgery. As before, inclusion criteria was sufficient renal function and B12 levels. Median (interquartile range) serum MMA levels in our control group was 140 (100–220) nmol/L, which is similar to serum MMA levels in the general population (14). Median serum MMA levels in patients with stage 4 NSCLC, however, was significantly higher: 260 (227–397) nmol/L (**Figure 1B**). We subsequently asked if MMA elevation seen in the serum of patients with Stage 4 disease was elevated compared to the *general* population. To obtain a comparator group of adults without lung cancer, we used the National Health and Nutrition Examination Survey (NHANES) database, which includes serum MMA measurements from a sample of 17,071 adults representative of the U.S. population. When we compared the serum MMA levels of our stage 4 NSCLC patients to those of NHANES participants, we found the median to be higher 260 (227–397) nmol/L vs 132 (101–178) nmol/L. To examine whether differences in MMA between our stage 4 NSCLC patients and NHANES participants could be explained by differences in the distributions of age, renal dysfunction, and vitamin B12 deficiency, we matched the cases 1:3 with cancer-free controls from NHANES based on age, vitamin B12 level, and serum creatinine. In a conditional logistic regression model, higher MMA levels remained associated with higher odds of having lung cancer (**Table S1**). These results suggest that the higher MMA levels observed in stage 4 NSCLC are most likely due to lung cancer and not confounded by age, ethnicity or comorbidities. Collectively, our data show that MMA is elevated in lung tumors and in the serum of patients with lung cancer, suggesting that propionate metabolism is dysregulated in NSCLC.

### MMAB is downregulated in lung cancer

MMA levels are controlled by six key proteins (9, 10, 20) (**Figure 2A**). If *MMAB, MMAA, MUT*, or *MCEE* were downregulated or inactivated by mutation, MMA levels accumulate. Conversely, overexpression of *PCCA* or *PCCB* can drive increased MMA levels by generating upstream metabolites that saturate propionate metabolism and lead to elevated MMA (20) (**Figure 2A**). We hypothesized that elevated serum and tumor MMA levels may be explained by downregulation of *MMAA, MMAB, MUT*, or *MCEE* or by overexpression of *PCCA* and *PCCB*. We analyzed the National Cancer Institute (NCI) Cancer Proteomic Atlas (21) and observed that MMAB protein levels were downregulated in both lung squamous and lung adenocarcinomas compared to resected adjacent normal tissue (**Figure 2B** and **Figure 2C**). In contrast, MMAA protein levels were elevated in tumors compared to adjacent normal tissue (**Supplement Fig 2A).** MUT was overexpressed in adenocarcinomas and under expressed in squamous cell carcinomas (**Supplement Fig 2B).** MCEE protein was downregulated in lung adenocarcinomas but elevated in squamous cell carcinoma (**Supplement Fig 2C)**. PCCA and PCCB were not overexpressed in tumors samples (**Supplement Fig 2D** and (**Supplement Fig 2E**). Taken together, our protein expression analysis revealed that MMAB was the only propionate metabolism regulatory gene that was consistently altered in a direction that would disrupt propionate metabolism and increase MMA levels in NSCLC.

**Figure 2.**
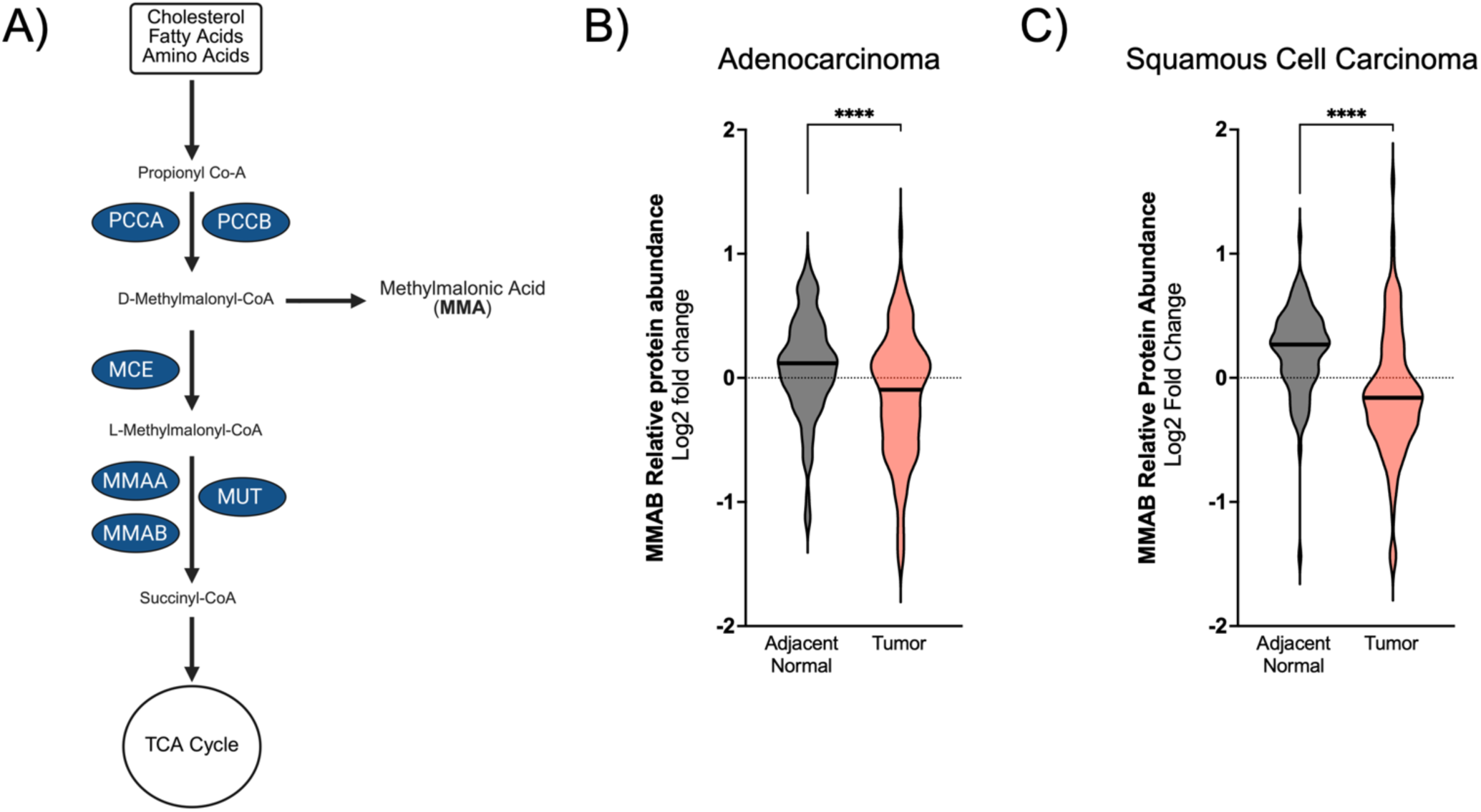
MMAB protein is downregulated in non-small cell lung cancer 1. **(A)** Schematic of propionate metabolism and the six key regulatory genes. **(B)** MMAB protein levels in tumor vs adjacent normal tissue in lung adenocarcinoma and squamous cell carcinoma from the NCI CPTAC proteomics data set. Paired Student’s t-test. ****p<0.0001

We subsequently asked if *MMAB* expression correlates with survival in NSCLC. We used KmPlot (22) to analyze bulk *MMAB* mRNA expression from patients with resected NSCLC. *MMAB* mRNA correlated with worse overall survival in lung adenocarcinomas (**Figure 3A**) but not squamous cell carcinomas (**Figure 3B**), indicating that MMAB downregulation may play an important role in lung adenocarcinoma.

**Figure 3.**
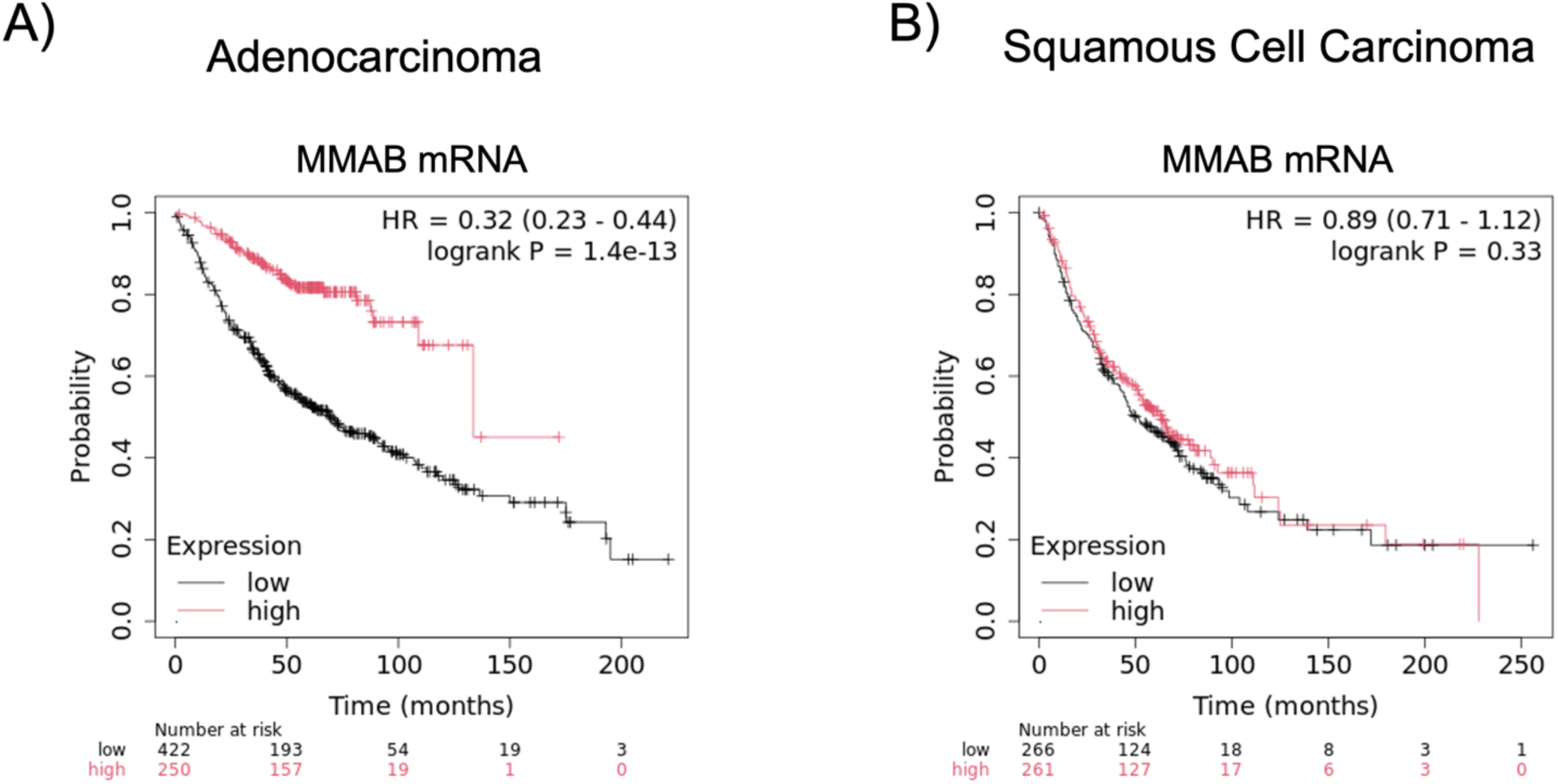
KMPLOT overall survival analysis of MMAB bulk mRNA expression from resected lung **(A)** adenocarcinoma and **(B)** squamous cell carcinoma.

### MMAB is downregulated in lung cancer cells

Analysis of bulk tumors showed that *MMAB* was downregulated in lung adenocarcinomas and correlates with overall survival. We next asked what cells express *MMAB* within the lung. To determine the expression pattern of *MMAB* at the single-cell level, we analyzed malignant tissue and adjacent normal lung samples from 37 patients who had early stage (Stage 1 and 2) lung adenocarcinoma (23). *MMAB* RNA expression was highest in the lung epithelial cells (**Figure 4A**) compared to stromal and immune cells (**Figure 4B**). Within the lung epithelium, *MMAB* expression was higher in AT2 and ciliated cells as compared to AT1 or basal cells (**Figure 4C).** *MMAB* was also significantly decreased in cancerous cells compared to AT2 cells, the presumed epithelial cell of origin for lung adenocarcinoma (**Figure 4D**)(24–26). Similar to our protein analysis, the changes in mRNA expression of *PCCA, PCCB, MCEE, MUT*, and *MMAA* did not support the hypothesis that their expression changes were disrupting propionate metabolism to drive increased MMA levels (**Figure S2**).

**Figure 4.**
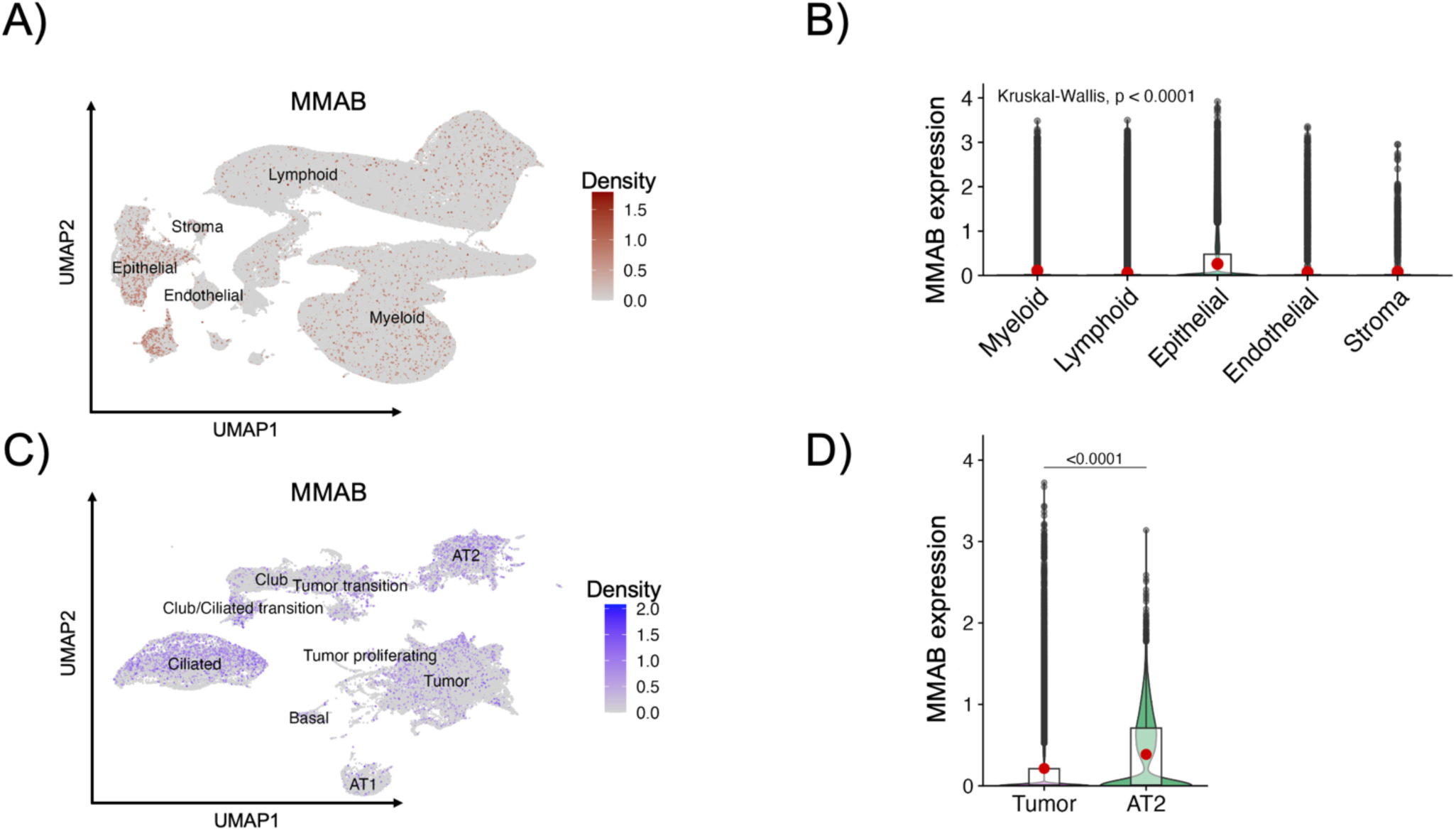
MMAB is expressed in the lung epithelium. **(A)** UMAP of single cell RNA-sequencing of 53,149 epithelial cells, 200,754 lymphoid cells, 193,379 myeloid cells, 7,906 endothelial cells, and 2,635 stroma cells from 37 patients with early-stage lung cancer. **(B)** Violin plot of MMAB expression within lung cell types. **(C)** UMAP of MMAB expression among lung epithelial and cancer cells. **(D)** Violin plot comparing MMAB expression in tumor cells vs. AT2 lung epithelial cells.

### Knockdown of MMAB increases MMA levels and activates EMT and Hypoxia pathways

Given *MMAB* is primarily expressed in lung epithelial cells and cancer cells, we asked if knocking down MMAB in lung cancer cell lines would increase MMA levels. Using shRNAs we knocked down MMAB in the A549 and H1975 lung cancer cell lines (**Figure 5A**) and measured MMA levels. MMAB knockdown increased intracellular MMA levels in both cell lines. To define the transcriptional effects of MMAB knockdown and MMA elevation in NSCLC cell lines, we conducted RNA-sequencing and analyzed genes that were commonly altered between both hairpins in each cell line. In the A549 cell line, 992 genes were significantly altered (**Figure 5B**) between shMMAB#1 and shMMAB#2. In the H1975 cell lines, there were 2,427 genes common between both hairpins. ENRICHR (27, 28) analysis revealed MMAB knockdown significantly enriched for EMT and hypoxia pathway activation (**Figure 5C**).

**Figure 5.**
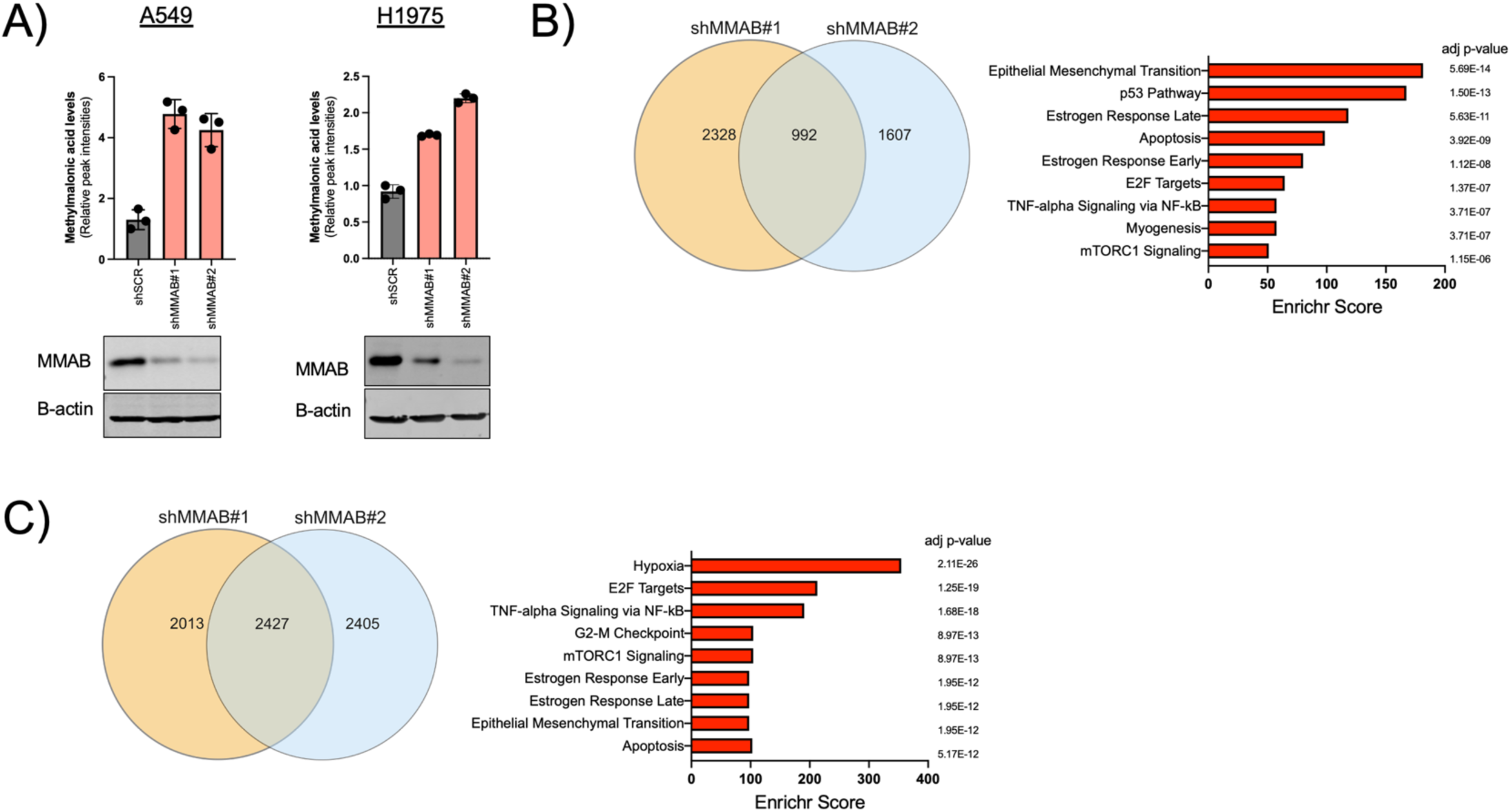
Knockdown of MMAB increases MMA and activates EMT and hypoxia gene signatures. **(A)** Targeted metabolomics of methylmalonic acid levels after MMAB knockdown in A549 and H1975 cells lines. **(B,C)** Pathway enrichment analysis of A549 and H1975 cell lines using genes commonly altered between two shRNAs against MMAB.

### MMAB is downregulated in persister cells

Hypoxia and EMT signaling are known to play important roles in promoting drug-tolerant persister (**DTP**) cells, or residual disease, in EGFR-mutated NSCLC by inhibiting apoptosis (29–32). We hypothesized that loss of MMAB and concomitant MMA elevation may promote DTP survival. To test this, we generated DTPs in three EGFR-mutant NSCLC cell lines by treating with osimertinib for 10 days (**Figure 6A**). MMAB was downregulated in all three cell lines (**Figure 6B**). To determine if MMA levels were elevated in DTPs, we performed targeted metabolomics for MMA in the H1975 and HCC827 cell lines and found that DTPs had significantly elevated MMA levels (**Figure 6C**). We subsequently asked whether MMAB was downregulated in clinical residual disease samples. We performed expression analysis of single-cell data published by Maynard *et al.* from patients with stage 4 EGFR-mutant NSCLC who were treated with an EGFR inhibitor (5). *MMAB* was downregulated in residual disease compared to treatment-naïve diseases (**Figure 6D**). Collectively, our data suggest that downregulation of MMAB and elevation of MMA may promote DTP survival.

**Figure 6.**
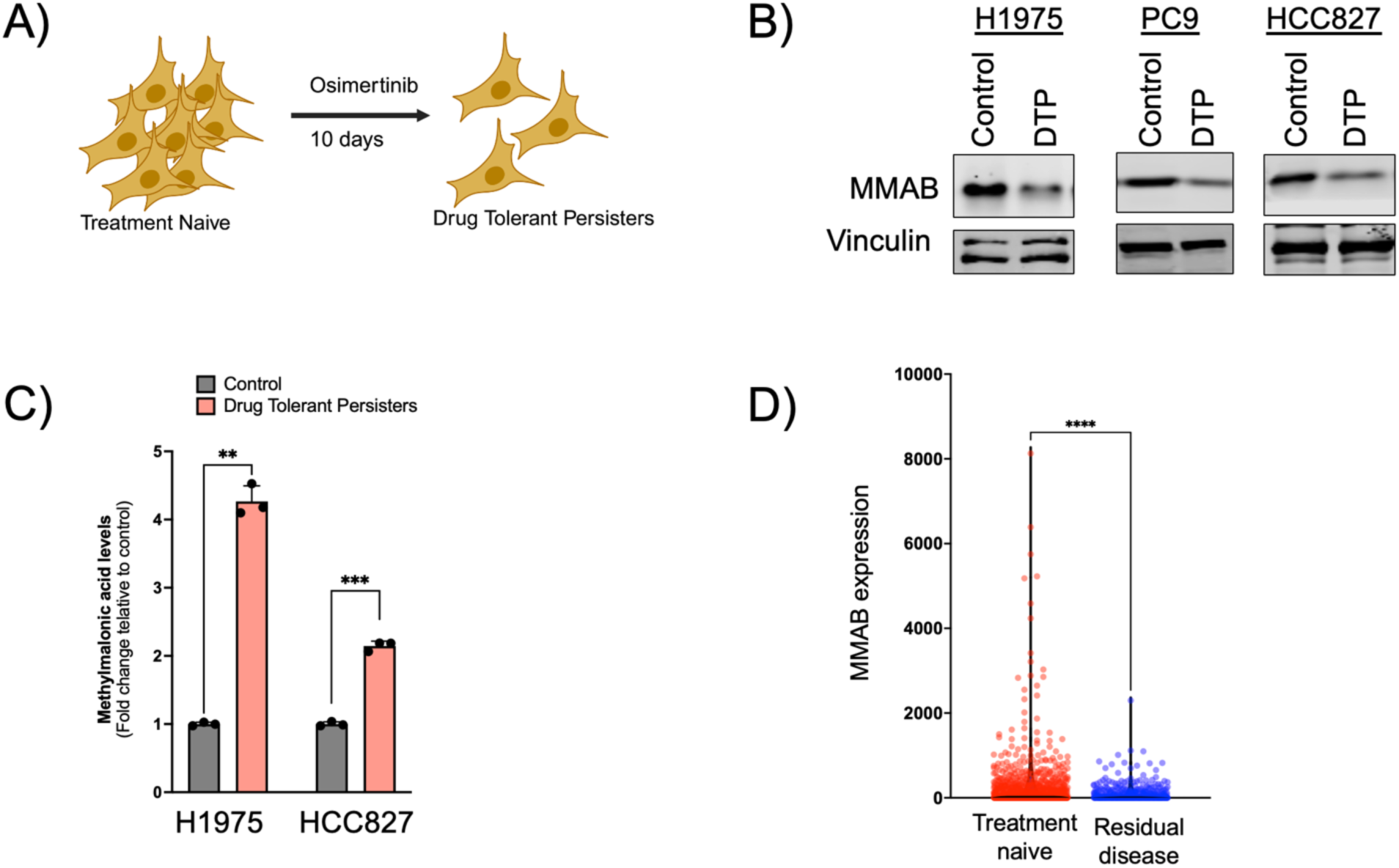
Propionate metabolism is dysregulated in drug-tolerant persister cells. **(A)** Schematic of generating drug-tolerant persister cells in H1975, PC9, and HCC827 EGFR-mutant cell lines. **(B)** MMAB protein expression from drug-tolerant persister cells. **(C)** Targeted metabolomics analyzing MMA levels in DTPs from H1975 and HCC827 cells. **(D)** Single-cell analysis of MMAB mRNA expression in treatment naïve and residual disease from patients who with Stage 4 EGFR-mutant NSCLC who were treated with targeted therapy.

### Overexpression of MMAB enhances osimertinib sensitivity

As MMAB is downregulated in DTPs, we asked if rescuing MMAB expression could increase osimertinib sensitivity. We transduced the H1975 and PC9 cells with an empty or V5-tagged MMAB construct. Overexpression of V5-MMAB mildly improved osimertinib sensitivity after 7 days (**Figure 7A** and **B**) of treatment. To test if MMAB expression plays a role in DTP survival, we utilized a colony formation assay and treated cells for 4 weeks. There was no significant difference in colony formation in control conditions, but V5-MMAB expressing cells were significantly more sensitive to long-term osimertinib treatment (**Figure 7C** and **D**).

**Figure 7.**
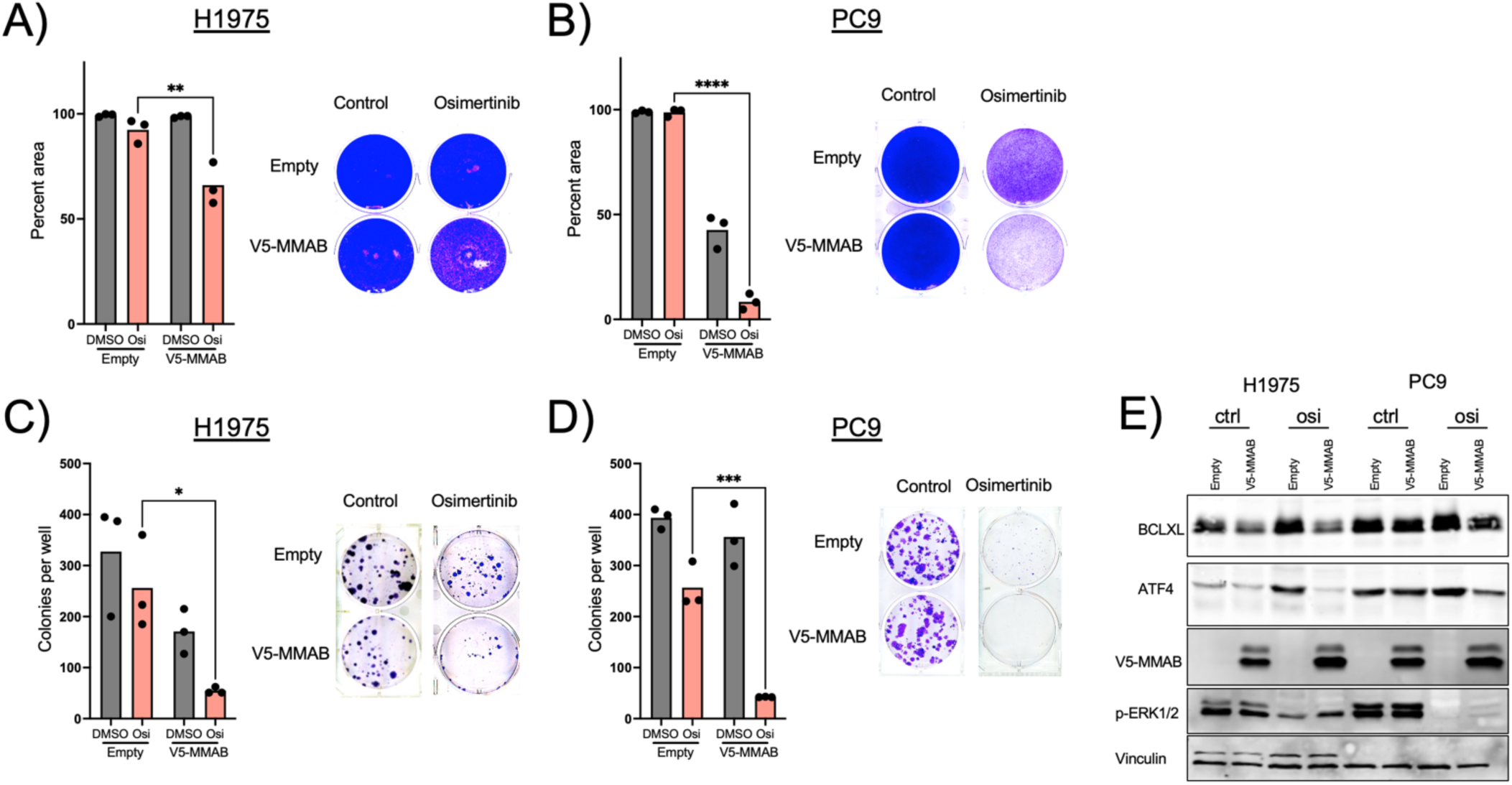
Restoring MMAB expression enhances osimertinib sensitivity. **(A-B)** H1975 and PC9 cells transduced with empty or V5-MMAB expressing vectors were treated with osimertinib for 7 days and stained with crystal violet. One-way ANOVA, **p<0.01, **** p<0.0001. **(C-D)** Colony formation assay of H1975 and PC9 cells after 4 weeks of Osimertinib treatment. One-way ANOVA, *p<0.05, *** p<0.001 **(E)** Western blot of H1975 and PC9 cells expressing empty or V5-MMAB vector treated with control or Osimertinib for 72 hours.

We have previously shown that elevated MMA levels activate TGFB signaling (18). We thus hypothesized that MMAB could be regulating TGFB to augment osimertinib sensitivity. To test this, we performed RNA-sequencing of V5-MMAB expressing H1975 cells. Ingenuity Pathway Analysis revealed that the top upstream network affected by V5-MMAB expression was the TGFB pathway (z-score −3.01 p 4.1e-36). Indeed, IPA analysis demonstrated that MMAB expression inhibited TGFB signaling. To confirm this, we tested whether ATF4 and BCL2L1, two TGFB targets that have been implicated in enabling DTP cell survival (33–35), were downregulated by MMAB expression at baseline and after 72 hours of osimertinib treatment (**Figure 7E**). In both the H1975 and PC9 cell line, we observed that MMAB expression blocked the induction of the anti-apoptotic genes ATF4 and BCL2L1.

## Discussion

Propionate metabolism is increasingly being implicated as playing an important role in human disease and aging, but its role in cancer remains incompletely understood. To determine its relevance in cancer, we asked if the pathway was disrupted in clinical specimen by measuring MMA levels in lung tumors and in the serum of patients with metastatic lung cancer. We also tested whether the expression of key genes involved in regulating propionate metabolism were altered in lung cancer. Collectively, we demonstrate that propionate metabolism is dysregulated in lung cancer, and that suppression of *MMAB*, an important regulator of the pathway, is necessary for DTP survival.

Previous work has shown that the disruption of propionate metabolism can promote drug resistance in cancer cell lines *in vitro* and metastases in mice *in vivo* (18–20). We have built on this work by showing the clinical relevance of propionate metabolism in lung cancer. Moreover, our findings show that *MMAB* may be the key enzyme that is altered in cancers that ultimately obstructs the pathway and leads to elevated MMA, which subsequently activates TGFB signaling and promotes therapy resistance. The precise mechanism by which MMA activates TGFB remains unclear. We speculate that MMA can signal either through G-protein coupled receptors (**GPCR**), similar to succinate and other TCA metabolites (36), or through chromatin remodeling as seen with some metabolites (37, 38). These are important outstanding questions for future studies to elucidate the role of propionate metabolism in human disease (9).

While *MMAB* has primarily been studied in the focused context of propionate metabolism and adenosylcobalamin regulation (39–41), recent studies have implicated *MMAB* in playing an important role in hepatocytes, including serving as a biomarker for toxicity and regulating cholesterol homeostasis (42, 43). Our work now shows that *MMAB* also plays a key role in DTP biology. We speculate that upon exposure to therapy *MMAB* is downregulated, leading to enriched intracellular MMA levels, which in turn activates TGFB signaling and EMT. Regulation of *MMAB* expression at the transcriptional and post-transcriptional levels is incompletely understood and is a key future question.

Eradicating residual disease in NSCLC is an area of great interest. Previous work has shown that the metabolic state of DTP cells is significantly different from proliferative, fully resistant cancer cells. Whereas fully proliferative cells employ anerobic glycolysis and oxidative phosphorylation, DTP cell metabolism is varied and complex. DTP cells rely on a mix of oxidative phosphorylation and aberrant fatty acid oxidation and redox regulation (44, 45). Our work establishes that propionate metabolism dysregulation is an important feature of DTP biology and suggests that restoring *MMAB* expression could be a therapeutic strategy.

## Methods

### Methylmalonic acid tumor measurement

Normal adjacent and tumor tissue samples were collected from a lung cancer biobank at Weill Cornell Medicine (IRB #1008011221). At the time of resection, normal adjacent and tumor tissue were snap frozen in liquid nitrogen. Approximately 10mg of tissue was collected and lysed in 80% ice-cold methanol. Targeted liquid chromatography tandem mass spectrometry (LC-MS/MS) was performed to detect a panel of polar metabolites according to previously published protocols (46). Samples were normalized by weight and analyzed using Ingenuity Pathway Analysis.

### Methylmalonic serum measurement

Serum from patients with NSCLC or a history of NSCLC were collected using an IRB approved protocol (IRB# 20-03021641) at Weill Cornell Medicine. Samples were sent de-identified to ARUP Laboratories (Salt Lake City, Utah) for measurement of MMA and vitamin B12.

#### NHANES Analysis

We downloaded data from the survey years 2001-2002, 2003-2004, 2011-2012, and 2013-2014, representing all data collection years with available MMA measurements, from the NHANES website (https://wwwn.cdc.gov/nchs/nhanes/default.aspx). For each respective year, we obtained MMA, vitamin B12, and serum creatinine levels from the Laboratory Data tables. The detailed methodology of measurement for each laboratory value is described on the NHANES website. We also obtained participant characteristics from Demographics Data and self-reported cancer history from Questionnaire Data. We calculated the median and interquartile range of MMA by 10-year age category for all participants and for the subgroup with vitamin B12 > 300 pg/mL and serum creatinine < 1.3 mg/dL to exclude participants with potential vitamin B12 deficiency or renal insufficiency. To examine whether serum MMA levels were higher in Stage 4 NSCLC patients compared to representative cancer-free individuals in NHANES, we created a dataset including all 24 cases of Stage 4 NSCLC matched 1:3 with controls from NHANES based on age, vitamin B12, and creatinine levels. We then performed conditional logistic regression with lung cancer as the outcome and MMA categorized as quartiles as the main predictor. To account for residual confounding, we adjusted our model for age, vitamin B12, and creatinine levels.

### MMAB expression analysis

For protein analysis, the NCI CPTAC database was queried using cprosite.ccr.cancer.gov. Data from paired non-malignant normal and tumor tissue was downloaded and analyzed in Graph Pad Prism using a paired t-test. For survival analysis, KmPlot was queried for each propionate metabolism gene at kmplot.com.

### Single cell RNA sequencing analysis

Single-cell RNA-sequencing data used in this study were generated previously by using the 10x Genomics Chromium platform and Illumina NovaSeq 6000 sequencing, and analyzed as described therein (23). Briefly, raw reads were aligned to the human transcriptome (GRCh38) using Cell Ranger (10x Genomics) with default parameters, followed by quality control to remove low-quality cells, doublets, and contaminating populations (47). The decontX algorithm was applied to mitigate ambient RNA contamination, and doublets were identified using DoubletFinder (48, 49). Processed data were normalized and integrated using the SCTransform workflow in Seurat with Harmony for batch correction (enabling identification of major cell lineages (epithelial, endothelial, stromal, lymphoid, and myeloid) based on canonical marker expression (50, 51). Each lineage was subsequently reclustered to refine annotations. Tumor epithelial cells were distinguished from normal epithelial cells by copy number variation analysis with InferCNV (https://github.com/broadinstitute/inferCNV), supported by aberrant marker expression and enrichment in tumor samples. P-values were calculated using the Kruskal–Wallis test and the Wilcoxon signed-rank test.

### Cell Culture

The H1975 and HCC827 cells were purchased from ATCC (CRL-5908 and CRL-2868). The PC9 cells were a gift from the laboratory of Dr. Harold Varmus. Cells were maintained in RPMI Media (Corning) with 10% FBS (Sigma) and penicillin-streptomycin (Gibco). MMAB shRNA constructs were purchased from Sigma (#TRCN0000083904 and #TRCN0000083906). A scrambled shRNA was used as a negative control (#SHC002). All shRNA constructs contained puromycin resistance and were selected for 3 days before use. A pDONR223 vector that contained V5-tagged MMAB was purchased from Horizon Discovery (#OSH 1770-202318739) and cloned into the pLX304 destination vector (Addgene #25890) using Gateway cloning. An pLX304 expression vector that did not contain a gene was used as an “empty” control. All cells transduced with a pLX304 vector were selected for using blasticidin for 7 days before use.

Bulk RNA sequencing was conducted on A549 and H1975 shMMAB cell lines according to prior methods (19). Briefly, after puromycin selection, cells were plated in 10cm plates in triplicate. RNA was then extracted 48 hours after plating using the Ambion PureLink RNA Minikit (Life Technologies) according to the manufacturer’s instructions and treated with DNAse 1 (Sigma). Library preparation and sequencing were performed at the Genomics Core at Weill Cornell Medicine.

#### Drug tolerant Persister Cells

Drug tolerant persister cells were generated by plating 300,000 cells into 10cm plates. Cells were then treated with osimertinib (Selleck) for 10 days. H1975 cells were treated with 1uM osimertinib, PC9 cells were treated with 2uM osimertinib, and HCC827 were treated with 20nM osimertinib.

### Osimertinib sensitivity assays

Approximately 300,000 H1975 or PC9 cells were plated in 6-well plates in triplicate and treated with 500nM or 1uM osimertinib, respectively. Media was refreshed every 48 hours. After 7 days, cells were washed with PBS and stained with crystal violet (Sigma C0775). For colony formation assays, 500 cells were plated into 6-well plates. After 24 hours, media was refreshed and osimertinib was added. Media was refreshed every 48-72 hours. At the conclusion of the experiment, media was aspirated, cells were washed with PBS and then incubated in crystal violet (Sigma) for 20 minutes at room temperature. Plates were then washed with distilled water and left to air-dry overnight. Plates were then scanned and DTP area was quantified using Image J.

#### Western Blotting

Cells were lysed in RIPA buffer (Cell Signaling Technology) containing phosphatase and protease inhibitors. All immunoblots were conducted on the LiCor Odyssey system. Primary antibodies were diluted in Odyssey blocking buffer with 0.1% Tween at a concentration of 1:1,000. Secondary antibodies were used at ∼1:10,000 dilutions. Proteins of interest were probed with the following primary antibodies overnight: V5 (Cell Signaling), Vinculin (Abcam), MMAB (Novus), ATF4 (Cell Signaling), BCL2L1 (Cell Signaling), p-ERK 1/2 (Cell Signaling), B-actin (Sigma).

## Statistics

For paired analyses, a paired Student’s t-test was used. For unpaired datasets, an unpaired Student’s t-test was performed. One-way ANOVA followed by post-hoc Sidak’s multiple comparisons test was conducted for comparisons among more than two groups. All statistics were performed in Graph Pad Prism.

## Study Approval

The studies with human tissue and blood were performed in accordance with approved protocols by Weill Cornell University Institutional Review Board and conducted in accordance with the principles of the Declaration of Helsinki. Informed consent was obtained from all participants.

## Data Availability

The datasets for all RNA-sequencing have been deposited at Array Express (Accession #E-MTAB-15737 and #E-MTAB-15738). Further inquires can be directed to the corresponding authors.

## Author Contributions

BP designed and performed experiments, analyzed data, and wrote the manuscript. LY acquired and analyzed data. RK performed experiments. ZL and MN acquired data and contributed to writing the manuscript. EEG designed experiments and analyzed data. YS and RH acquired and analyzed data. AS and NA acquired data and analyzed data. JB designed experiments, analyzed data, and contributed to writing the manuscript.

## Acknowledgments

We thank the members of the Blenis laboratory for their helpful comments and feedback, and we thank the Cornell Institute of Biotechnology Proteomics and Metabolomics core for their services. We also thank Dr. Eunji Choi for technical assistance.

---

**Figure S1.**
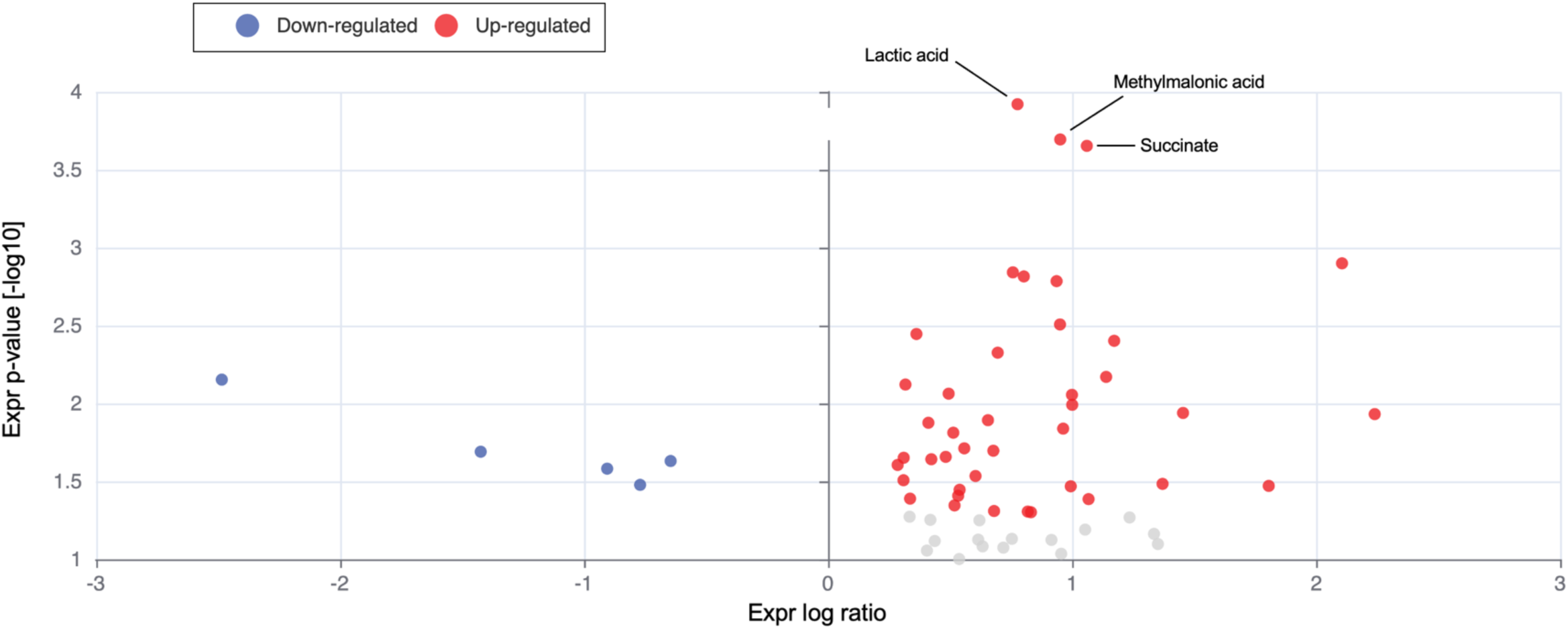
Volcano plot of metabolites from tumor compared to non-malignant normal lung tissue. The x-axis represents log2 fold change and the y-axis represents the −log10 p-value with higher values indicating greater statistical significance. Plot generated by Ingenuity Pathway Analysis.

**Supplementary Table 1.** Serum MMA is elevated in patients with metastatic lung cancer compared to serum from the general population. Odds ratios for having lung cancer depending on serum MMA levels after adjusting for age, sex, vitamin B12 deficiency, and kidney dysfunction.

| **MMA Quartile** | **Odds ratio** | **95% CI** | **p-value** |
| --- | --- | --- | --- |
| 1 | - | - |  |
| 2 | 1.45 | 0.25, 7.74 | 0.7 |
| 3 | 3.46 | 0.63, 19.0 | 0.2 |
| 4 | 5.69 | 1.17, 27.7 | 0.031 |

**Figure S1.**
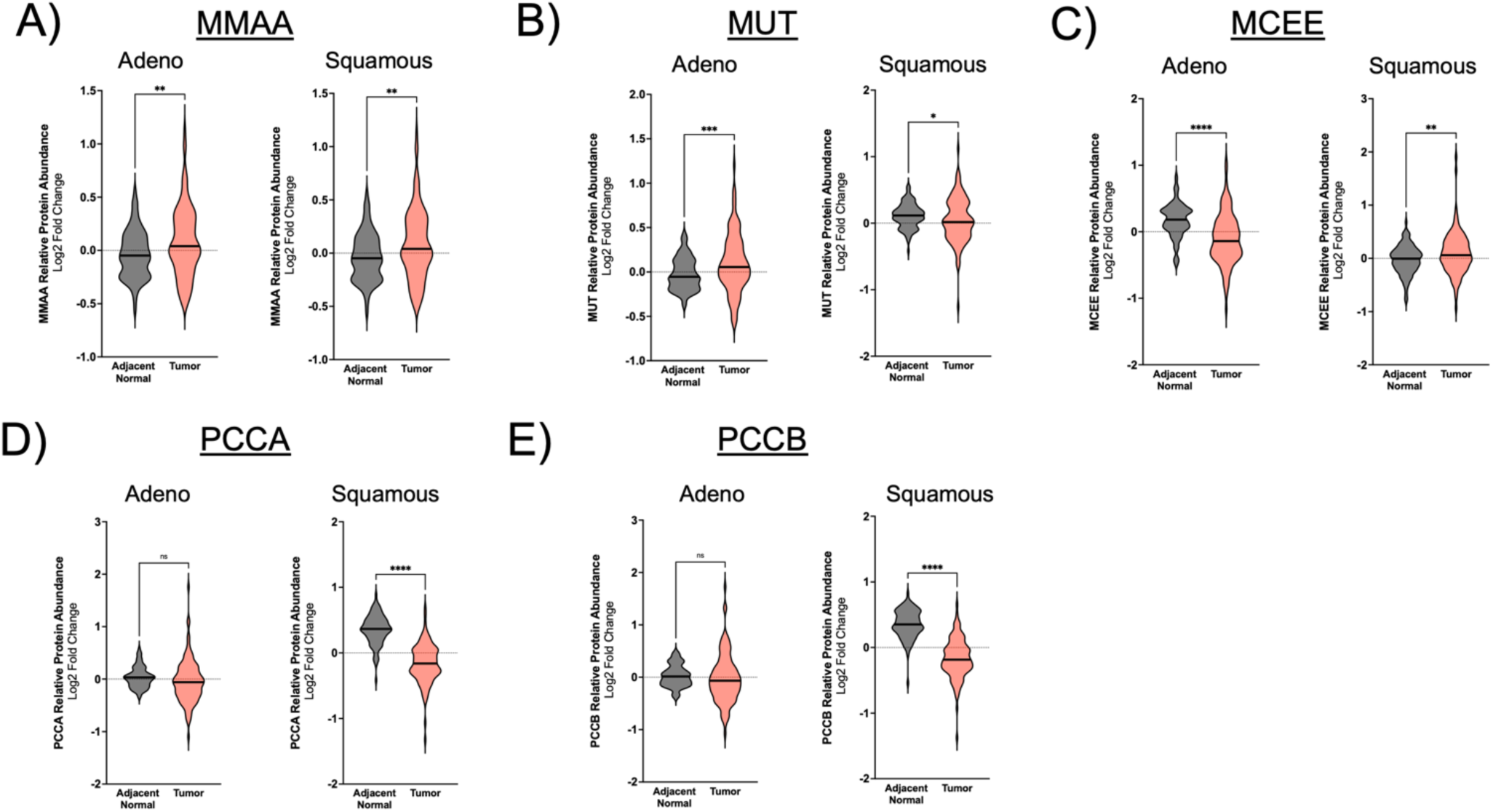
**(A)** MMAA **(B)** MUT **(C)** MCEE **(D)** PCCA and **(E)** PCCB protein levels in tumor vs adjacent normal tissue in lung adenocarcinoma and squamous cell carcinoma from the NCI CPTAC proteomics data set. Paired Student’s t-test. *p<0.05, **p<0.01, ****p<0.001

**Figure S2.**
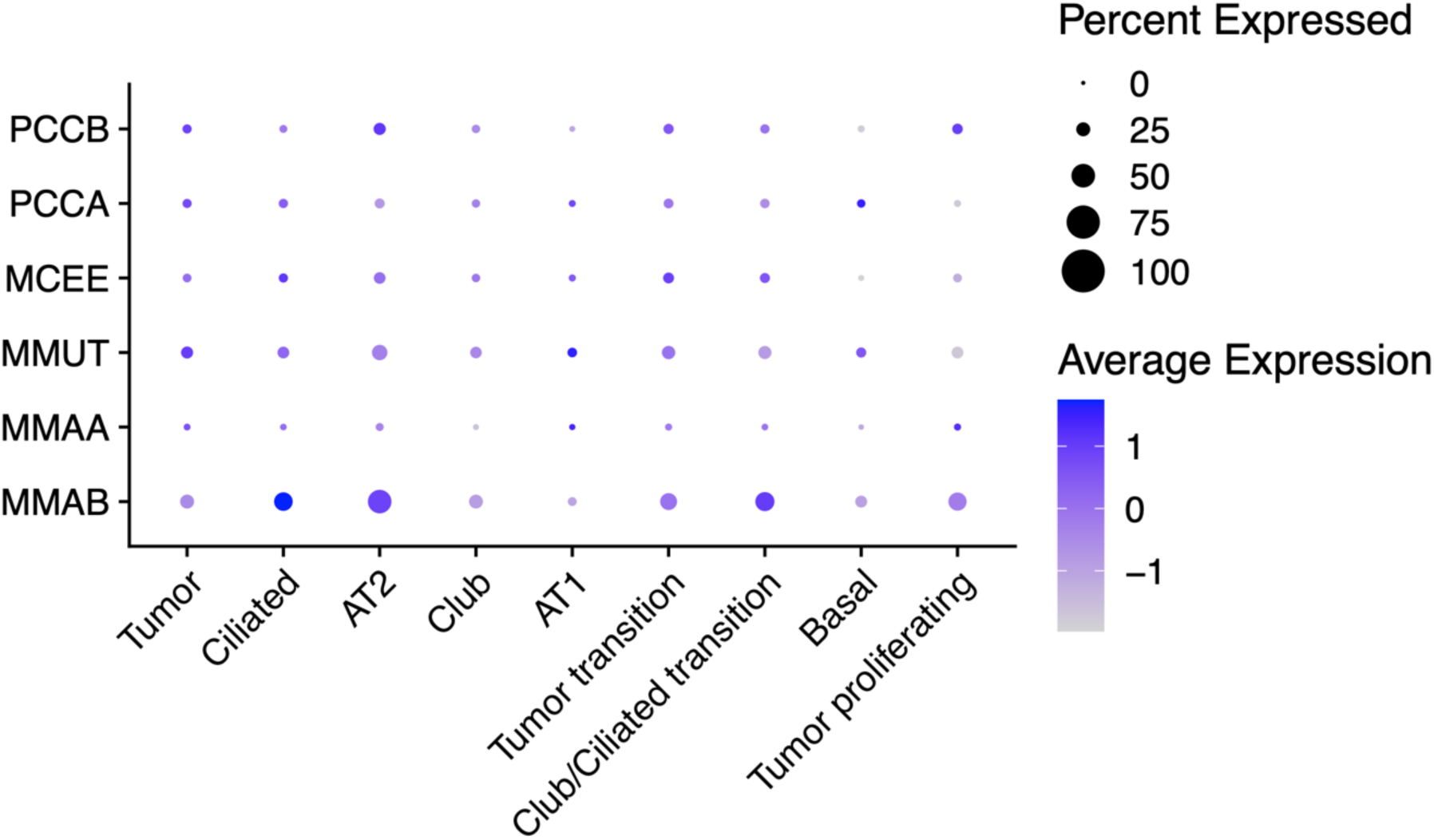
Dot plot of differential single-cell expression of propionate metabolism genes lung cancer

